# Efficient amplicon panels for evaluating hybridization and genetic structure in the *Schistosoma haematobium* species group

**DOI:** 10.1101/2025.07.18.665602

**Authors:** Roy N Platt, Grace A Arya, Bonnie L Webster, Aidan M Emery, Timothy JC Anderson

**Author notes:** Correspondence to RN Platt II or TJC Anderson.

## Abstract

Genetic markers for detecting hybridization and measuring population genetic parameters must be informative and cost-effective. Most population genetic studies within the *Schistosoma haematobium* species group rely on either a two-marker system consisting of the mitochondrial, <u>c</u>ytochrome <u>ox</u>idase <u>1</u> (*cox*1) and the nuclear internal transcribed <u>s</u>pacer (ITS) markers or, at the other extreme, millions of <u>s</u>ingle <u>n</u>ucleotide <u>v</u>ariants (SNVs) from whole genome/exome sequencing. *cox*1 and ITS studies contain minimal population genetic information but whole genome sequencing is cost-prohibitive. We examined ∼38 million previously published, whole genome SNVs genotyped in 162 *S. haematobium* and *S. bovis* sampled across Africa. We compared population genetic parameters from 4,000 panels of 10-100,000 randomly sampled SNVs to results from the whole genome dataset to test the resolution of reduced representation sequencing in schistosomes. We found that panels of 500 SNVs captured >99% of the population genetic information contained in the whole genome dataset by using Procrustes transformed principal component analyses and ancestry estimates (*r²* = 0.85). Additionally, the costs of genotyping parasites with an amplicon panel is two to three times less than whole genome sequencing. Our results show that moderately-sized amplicon panels targeting random SNVs provide an efficient approach to large scale, field-based schistosome surveillance.

## Background

Many schistosome species are capable of hybridizing under laboratory conditions (1) and the opportunity for hybridization in the field is a particular concern (2, 3). Significant efforts have been made to monitor the extent of hybridization in the field, with a strong focus on interbreeding between the livestock parasite, *S. bovis,* and the human parasite, *S. haematobium*. The discordance between mitochondrial <u>c</u>ytochrome <u>ox</u>idase subunit I (*cox*1) and nuclear ribosomal internal transcribed <u>s</u>pacer (ITS) genetic markers has been used to identify putative hybrid schistosomes collected from humans in many African countries (4). Despite the reliance on this system, there are reasons to believe that hybrid parasites cannot be accurately identified using just two markers.

A central problem is that hybrid parasites that contain heterozygous alleles at ribosomal ITS loci could reflect F1 hybrids, or be the result of shared ancestral polymorphisms and lineage sorting (5). Equally well, parasites resulting from historical hybridization events that occurred hundreds or thousands of generations ago may retain heterozygous ITS alleles further complicating hybrid identification. To provide a parallel example, many humans contain low levels of Neanderthal or Denisovan DNA, most of which are present in heterozygous state (6). These heterozygous archaic human alleles obviously do not indicate that F1 human-Neanderthal hybrids but instead reflect events that occurred ∼50,000 years ago.

The presence of discordant mitochondria, represented by the *cox*1 gene, can be caused by hybridization, incomplete lineage sorting as well as introgression. Due to their haploid and non-recombining nature, multiple discrete mitochondrial haplotypes can circulate within a population. When populations sub-divide, these haplotypes may be randomly distributed in the daughter lineages. As a result, the phylogenetic history of the mitochondria and alleles in the nuclear genome may differ substantially and give the appearance of gene flow between non-sister taxa (7). In other cases, mitochondrial discordance can be the direct result of historical gene flow between species which has been well documented in (8), flycatchers (9), skuas (10), chipmunks (11), *Daphnia* (12), and shrews (13), among other species.

As a potential solution, whole genome studies can generate millions of <u>s</u>ingle <u>n</u>ucleotide <u>v</u>ariants (SNVs) and provide higher resolution than *cox*1 and ITS markers. Unfortunately, the data is expensive to generate which limits the number of samples that can be examined in a study. To date, only a few hundred field-collected *S. haematobium* and *S*. *bovis* parasites have been examined with genome or exome SNVs compared to the thousands of samples included in *cox*1 and ITS studies. These whole genome or exome studies have not yet identified recent hybrids (F1 or F2), but are underpowered for detecting rare contemporary hybridization events (14–17). Hence while the precision of whole genome sequencing is desirable, such studies are cost prohibitive, and difficult to scale.

Several techniques have been used to generate large numbers of genome-wide markers in a high-throughput and cost-effective way but each has limitations. Microsatellites can generate dozens of distributed markers with multiplex PCRs and these results have confirmed the reproductive isolation between *S. haematobium* and *S. bovis* in Senegal (15). However, scoring microsatellites is not straightforward and can vary between labs (18). Reduced representation sequencing methods (19), like restriction-site <u>a</u>ssociated <u>D</u>NA <u>seq</u>uencing (RADseq), have shown that *S. haematobium* hybridizes with *S. guineensis* (20), but reproducibility between RADseq studies can be an issue in general (21). Targeted sequence capture which uses biotinylated probes to hybridize and enrich pre-determined loci have been used to sequence entire exomes in schistosomes (17, 22–24). The probes themselves are capable of capturing moderately diverged sequences making them particularly useful in capturing homologous loci in related taxa (25), but the initial cost of developing the probes and cost per reaction can represent a hurdle. Amplicon panels can be generated to target multiple markers simultaneously using a two-step multiplexed PCR reaction (26). In the first step ancestry-informative markers (27), drug resistance markers (28), small indels (29) or other combinations of variants (30) can be targeted with locus specific primers. Amplicon panels that target moderate numbers of SNVs have been show to resolve population structure for a range of taxa with dozens to hundreds of markers (for examples see 31, 32, 33) and primer sequences can be shared between labs potentially solving the problem of reproducibility.

A clear need exists for a high-throughput cost effective method for genotyping schistosomes that is robust and reproducible. This study was designed to determine how many SNVs are required for accurate assessment of hybridization and population structure in schistosome populations using amplicon panels. To achieve this, we down sampled SNVs from a published dataset (14) comprising 38 million SNVs in 141 *S. haematobium* and 21 *S. bovis* parasites collected across Africa. The genome wide dataset clearly defines three populations: *S. bovis*, northern *S. haematobium* and southern *S. haematobium*. We compare panels of various sizes to determine the minimum number of markers necessary to maintain high accuracy in ancestry estimation for investigating hybridization and population structure in schistosomes. Our findings demonstrate that panels containing several hundred, genome-wide markers can perform as well as whole genome genotyping. Adoption of such panels has the potential to transform molecular surveillance in endemic areas, enabling more accurate detection of hybrids and better informing control strategies, while maintaining scalability and affordability.

## Methods

### Down-sampling

All sequence data used in this study were obtained from public sources and previously examined in Platt *et al*. (14). The SNVs are available as a single file in the variant call format (VCF) at https://doi.org/10.5061/dryad.xgxd254sk. The dataset includes 38,198,427 SNVs genotyped for 162 *S. haematobium* (n=141) or *S. bovis* (n=21) parasites. *S. haematobium* individuals were further subcategorized into two populations that broadly coincide with north (n=82) and south (n=59) Africa based on previous analyses (14). We applied additional filters to retain autosomal variants at moderate to high frequencies (>5%) using VCFtools v0.1.16 (34). Then we used PLINK v 2.00a5.12 to prune linked variants in sliding windows of 25 SNVs, shifting the window by 5 SNVs at a time, and removing variants with pairwise linkage disequilibrium, measured as *r*²>0.2.

We randomly subsampled SNVs from the filtered dataset in 38 bins, ranging in size from 10-100,000 SNVs. This was replicated 100 times by randomly sampling SNVs without replacement and repeating the population genetic analyses (described below). Random SNVs were selected using Python’s v3.11.11 random.sample() function, with the random seed initialized using NumPy’s v2.1.3 randint() function (35). A VCF file was created for each SNV subset using VCFtools v0.1.16 (34).

### Population genetic comparisons using down-sampled datasets

We compared the complete dataset, with each down-sampled dataset using several metrics:

(i) *Principal component analysis (PCA)*: This analysis was performed using PLINK v 2.00a5.12 (36). Each subset PCA was compared to a reference PCA, as generated from all filtered SNVs, using a Procrustes transformation (37) via the scipy.spatial.procrustes() function v1.15.2 (38). Disparity (*M²*) is a unitless metric that minimizes the sum of the squared distances between data points (individuals) in the transformed PCAs. Procrustes disparity (*M²)* can be converted to similarity (𝑡_0_) with 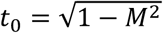. In this case it is used to assess differences in population structure as it is ascertained by PCA with all of the SNVs versus a subset.
(ii) *Fixation index (FST):* We calculated the global, weighted, Weir-Cockerham FST for each subsampled set of SNVs. Individuals in the data set were assigned to three possible populations including *S. bovis* and either northern or southern *S. haematobium* based on the designations specified in Platt *et al*. (14). We measured FST between populations with VCFtools v0.1.16 (34).
(iii) *Ancestry estimation*: We estimated ancestry for each individual using panels with 10-100,000 SNVs from a randomly chosen set of replicates. We estimated ancestry for each panel and included two populations (*k*) and 1,000 cross-validation replicates with ADMIXTURE v1.3.0 (39). We selected *k*=2 because it corresponds with the two species in the dataset; *S. haematobium* and *S. bovis*. Next, we compared the non–*S. haematobium* ancestry proportion (*q*) between the subsampled and full datasets using the linear regression linregress() function from SciPy v1.15.2 (38). The linear regression was restricted to *S. haematobium* individuals based on the species assignments provided by Platt *et al*. (14). We calculated the slope, intercept, correlation (*r²*), and p-value to assess the strength and significance of the association between the two sets of ancestry estimates.

### Cost estimates

We estimated the costs of genotyping 1,000 parasites with whole genome sequencing, and an amplicon panel with 500 loci. We assumed that each sample started from a single miracidium stored on an FTA card. DNA is extracted from the card and amplified simultaneously using whole genome amplification (23). We targeted 20x coverage for whole genome samples based on a 400.3 Mb (40) genome (8.0 Gb / parasite) and 100x coverage per sample per locus for amplicon panels (7.5 Mb / parasite). Costs for reagents were taken directly from manufacturer websites and do not include unpublished discounts. When reaction sizes were not divisible by 100 samples, we included costs for partial kits.

### Reproducibility

Analyses were conducted on compute nodes with 192 cores and 1 TB of memory at the Texas Biomedical Research Institute’s high-performance computing cluster. We managed all software environments with Conda v24.11.3 and open source repositories at CondaForge MiniForge and BioConda. All code necessary to reproduce the analyses are stored in a Jupyter Lab v4.3.5 notebooks and environmental recipe files at GitHub (https://github.com/nealplatt/schisto_aim_panel) and permanently archived on Zenodo at (https://doi.org/10.5281/zenodo.15236800).

## Results

The Platt *et al*. (14) dataset contained 162 *S. haematobium* and *S. bovis* samples genotyped at 38,198,427 SNVs. We removed 7,860,528 variants from the mitochondrial genome and sex chromosomes and another 24,364,917 that were at low frequency (<5%). We then removed an additional 5,591,481 variants that displayed high linkage disequilibrium. After filtering, the >38 million SNVs from the Platt *et al*. (14) dataset were reduced to 381,501 moderate-to-high frequency, unlinked, autosomal variants.

We randomly down sampled the variants into datasets with 10-100,000 SNVs and replicated each SNV size category 100 times to understand how many SNVs are needed to replicate the results from the full, genome-wide dataset. In total we generated 3,700 random combinations of SNVs. For each set of SNV replicates we compared a PCA to one generated from the genome-wide SNVs in the filtered dataset (Table 1). Similarity between the Procrustes transformed PCAs is correlated with the number of markers examined (Figure 1); with 10 markers similarity between the PCAs was low (mean 0.755; range = 0.422-0.886). By 500 SNVs mean similarity between all replicated and whole genome datasets was high (*t_o_*≥0.99) with an interquartile range <0.001. At 100,000 markers the PCAs of each replicated dataset was effectively identical to the whole genome dataset (*t_o_* > 0.999; Figure 2).

**Figure 1.**
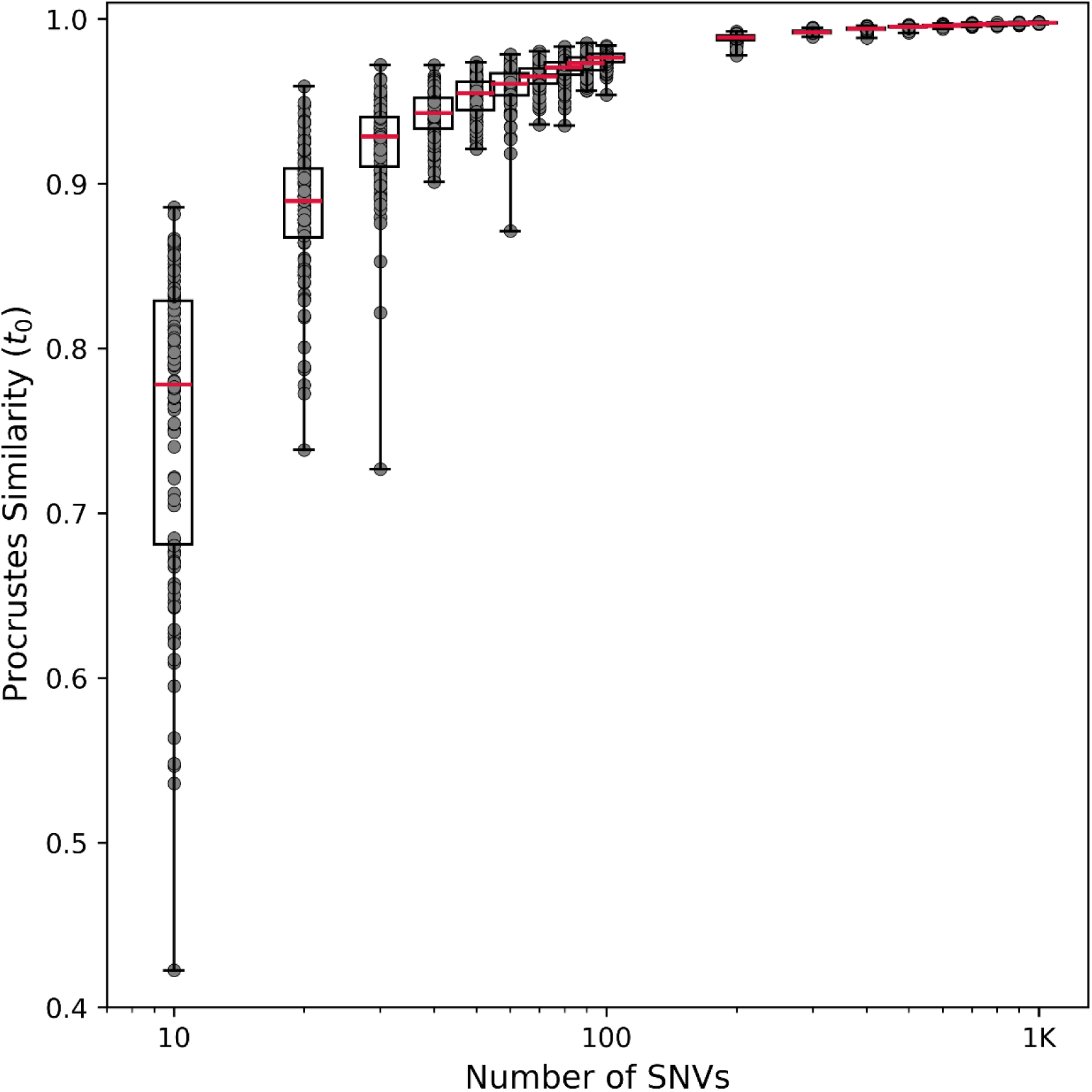
Similarity between randomly sampled SNV panels and a whole genome dataset. (A). Procrustes similarity (*t_0_*) between panels of random SNVs and the whole genome dataset were calculated across >100 replicate PCAs for panels ranging in size from 10-1,000 SNVs. SNV datasets >1,000 SNVs aren’t shown but show *t_0_*<0.999 in all replicates. Red lines indicate mean values, box plots cover the interquartile range from 25%-75% of values. Whiskers indicate the full range of values. All *t_0_* values are shown with grey markers. The source data shown in this figure is available in the supplemental materials.

**Figure 2.**
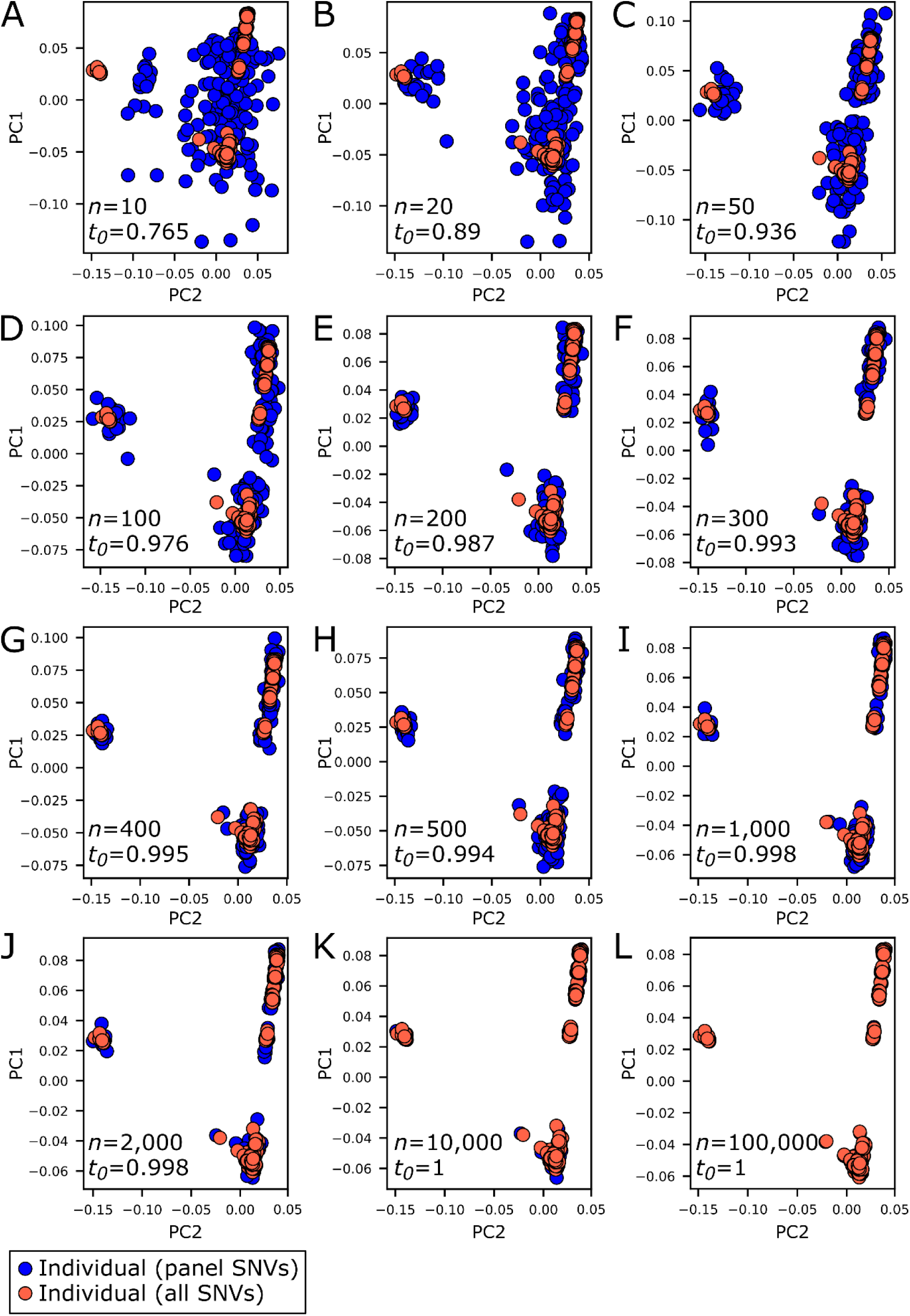
<u>P</u>rincipal <u>c</u>omponent <u>a</u>nalyses (PCA) converge with increase numbers of SNVs. PCAs were generated from all filtered variants (n = 381,501 SNVs; red markers) and randomly selected variants (blue markers). Procrustes transformations were used to measure similarity (*t_0_*) between PCAs generated from all and a random subset of variants. We examined SNVs panels containing 10 - 100,00. Results are shown for (A) 10, (B) 20, (C) 50, (D) 100 (E) 200, (F) 300, (G) 400, (H) 500, (I) 1,000, (J) 2,000, (K) 10,000, and (L) 100,000 SNVs at which point the similarity between the panel and full data sets are effectively identical. The source data shown in this figure is available in the supplemental materials.

**Table 1.** Procrustes similarity (t0) measured between SNV panels and the whole genome dataset across multiple replicates.

| # SNVs | Mean $t_0$ | Med. $t_0$ | Min $t_0$ | Max $t_0$ | $t_0$ IQR* | Successful Replicates |
| --- | --- | --- | --- | --- | --- | --- |
| 10 | 0.755 | 0.778 | 0.422 | 0.886 | 0.148 | 100 |
| 20 | 0.883 | 0.890 | 0.738 | 0.959 | 0.042 | 100 |
| 30 | 0.923 | 0.929 | 0.727 | 0.972 | 0.030 | 100 |
| 40 | 0.942 | 0.943 | 0.901 | 0.972 | 0.019 | 100 |
| 50 | 0.953 | 0.955 | 0.921 | 0.974 | 0.017 | 100 |
| 60 | 0.958 | 0.961 | 0.871 | 0.978 | 0.013 | 100 |
| 70 | 0.965 | 0.965 | 0.936 | 0.980 | 0.009 | 100 |
| 80 | 0.969 | 0.971 | 0.935 | 0.983 | 0.008 | 100 |
| 90 | 0.972 | 0.973 | 0.956 | 0.985 | 0.008 | 100 |
| 100 | 0.976 | 0.977 | 0.954 | 0.984 | 0.005 | 100 |
| 200 | 0.988 | 0.989 | 0.978 | 0.992 | 0.003 | 100 |
| 300 | 0.992 | 0.992 | 0.989 | 0.995 | 0.001 | 100 |
| 400 | 0.994 | 0.994 | 0.988 | 0.996 | 0.001 | 100 |
| 500 | 0.995 | 0.995 | 0.992 | 0.997 | 0.001 | 100 |
| 600 | 0.996 | 0.996 | 0.994 | 0.997 | 0.001 | 100 |
| 700 | 0.997 | 0.997 | 0.995 | 0.998 | 0.001 | 100 |
| 800 | 0.997 | 0.997 | 0.996 | 0.998 | 0.001 | 100 |
| 900 | 0.997 | 0.997 | 0.996 | 0.998 | 0.000 | 100 |
| 1,000 | 0.998 | 0.998 | 0.997 | 0.998 | 0.000 | 100 |
| 2,000 | 0.999 | 0.999 | 0.998 | 0.999 | 0.000 | 100 |
| 3,000 | 0.999 | 0.999 | 0.999 | 0.999 | 0.000 | 100 |
| 4,000 | 0.999 | 0.999 | 0.999 | 1.000 | 0.000 | 100 |
| 5,000 | 1.000 | 1.000 | 0.999 | 1.000 | 0.000 | 100 |
| 6,000 | 1.000 | 1.000 | 0.999 | 1.000 | 0.000 | 100 |
| 7,000 | 1.000 | 1.000 | 1.000 | 1.000 | 0.000 | 100 |
| 8,000 | 1.000 | 1.000 | 1.000 | 1.000 | 0.000 | 100 |
| 9,000 | 1.000 | 1.000 | 1.000 | 1.000 | 0.000 | 100 |
| 10,000 | 1.000 | 1.000 | 1.000 | 1.000 | 0.000 | 100 |
| 20,000 | 1.000 | 1.000 | 1.000 | 1.000 | 0.000 | 100 |
| 30,000 | 1.000 | 1.000 | 1.000 | 1.000 | 0.000 | 100 |
| 40,000 | 1.000 | 1.000 | 1.000 | 1.000 | 0.000 | 100 |
| 50,000 | 1.000 | 1.000 | 1.000 | 1.000 | 0.000 | 100 |
| 60,000 | 1.000 | 1.000 | 1.000 | 1.000 | 0.000 | 100 |
| 70,000 | 1.000 | 1.000 | 1.000 | 1.000 | 0.000 | 100 |
| 80,000 | 1.000 | 1.000 | 1.000 | 1.000 | 0.000 | 100 |
| 90,000 | 1.000 | 1.000 | 1.000 | 1.000 | 0.000 | 100 |
| 100,000 | 1.000 | 1.000 | 1.000 | 1.000 | 0.000 | 100 |

\*IQR = interquartile range

We examined changes in ancestry assignment (*q*) using Admixture and assuming 2 populations (*k*). Specifically, we compared *S. bovis* introgression in *S. haematobium* parasites by comparing *q* from SNV panels and the whole genome dataset (Figure 3). *S. bovis* ancestry estimates from whole genome data and 10 random SNVs was weakly and negatively correlated (Table 2, Figure 3). In this case, the ancestry component that is highest in *S. bovis* is also highest in southern *S. haematobium*. This result demonstrates the difficulty in ancestry assignment with underpowered datasets. Whole genome and panel estimates were highly correlated with 400 (*r^2^*=0.77) to 500 (*r^2^*=0.86) SNVs. The panel of 500 SNVs underestimated the amount of introgressed *S. bovis* ancestry by (mean) 0.4% but some samples were affected by as much as - 10.3% to 5%. Interestingly the mean bias is comparable in the 400 SNV panel (-0.6%) but the range of bias is smaller from -7.5% to 6%. This is likely an artifact of randomly selecting SNVs and would decrease with the number of markers. Once 1,000 SNVs were sampled all correlations were greater than 0.9 and the bias range is less than 8%.

**Figure 3.**
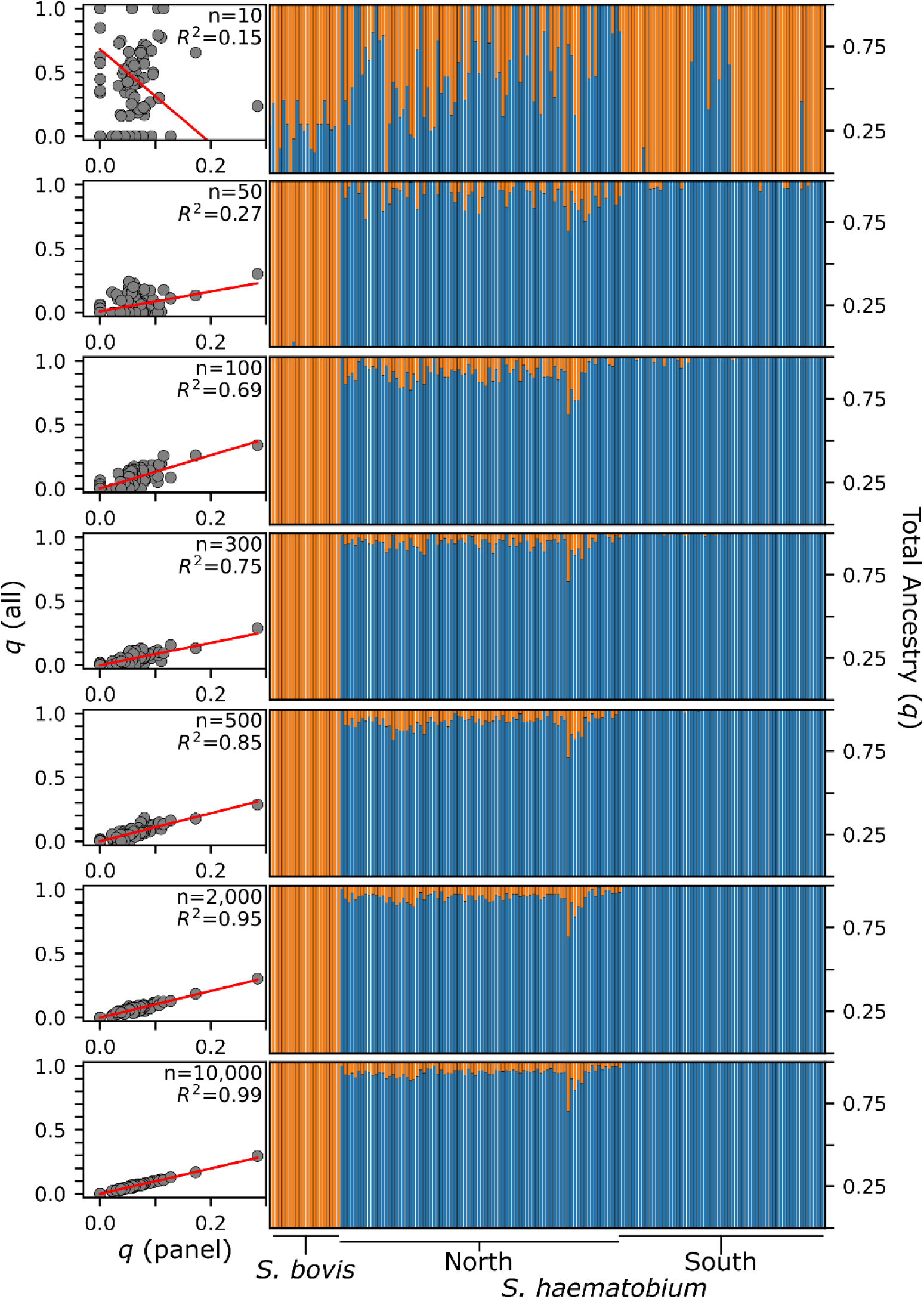
Comparing ancestry estimates SNV panels and whole genome data. (A). *S. bovis* introgressed alleles are present in many *S. haematobium* individuals. We compared the percent minor parent ancestry (*q*) from SNV panels to estimates made from whole genome data. The two are positively correlated once 50 SNVs are sampled though the strength of the correlation continues to increase. The scatter plots only show *S. haematobium* individuals. These results show that the panels of SNVs can accurately replicate a set of whole genome SNVs with a few hundred to thousand markers. The source data shown in this figure is available in the supplemental materials.

**Table 2.** Correlation among *S. haematobium* and *S. bovis* ancestry estimates from SNV panels and a whole genome dataset.

| Panel Size | $r^2$ | Slope | $p$-value | SE | Ancestry Bias | | | |
| --- | --- | --- | --- | --- | --- | --- | --- | --- |
|  |  |  |  |  | Skew | Mean | Min | Max |
| 10 | 0.153 | -3.67 | 1.60E-06 | 0.73 | 0.00 | -49.6% | -100.0% | 12.8% |
| 20 | 0.318 | 1.48 | 3.28E-13 | 0.18 | -1.95 | -2.7% | -45.0% | 9.5% |
| 30 | 0.536 | -4.64 | 6.06E-25 | 0.37 | 0.05 | -64.5% | -100.0% | -10.8% |
| 40 | 0.253 | 0.61 | 2.15E-10 | 0.09 | -0.63 | 0.9% | -16.0% | 10.5% |
| 50 | 0.272 | 0.77 | 3.36E-11 | 0.11 | -0.81 | -0.1% | -19.0% | 10.5% |
| 100 | 0.691 | 1.30 | 2.87E-37 | 0.07 | -0.66 | -1.5% | -14.3% | 7.4% |
| 200 | 0.637 | 1.23 | 2.15E-32 | 0.08 | -1.02 | -2.0% | -16.9% | 7.5% |
| 300 | 0.754 | 0.87 | 4.02E-44 | 0.04 | 0.27 | 0.6% | -5.6% | 8.1% |
| 400 | 0.774 | 1.03 | 9.34E-47 | 0.05 | -0.36 | -0.6% | -7.5% | 6.2% |
| 500 | 0.851 | 1.09 | 2.17E-59 | 0.04 | -1.10 | -0.4% | -10.3% | 5.3% |
| 1,000 | 0.921 | 1.07 | 1.46E-78 | 0.03 | -0.78 | -0.5% | -4.8% | 2.9% |
| 2,000 | 0.954 | 1.04 | 3.78E-95 | 0.02 | -0.04 | -0.1% | -3.6% | 3.0% |
| 10,000 | 0.992 | 0.99 | 8.87E-149 | 0.01 | 0.28 | 0.1% | -1.0% | 1.3% |
| 50,000 | 0.999 | 0.99 | 7.17E-209 | 0.00 | 0.43 | 0.1% | -0.4% | 0.5% |
| 100,000 | 0.999 | 1.01 | 1.33E-218 | 0.00 | -0.77 | 0.0% | -0.6% | 0.3% |

We measured the global, weighted, Weir-Cockerham FST for each replicate. Overall, the mean FST values were relatively robust to changes in panel size (Figure 4). For example, FST from all SNVs was 0.429 and the mean FST with 10 SNVs was 0.407 measured across 100 replicates. However, the variance in FST decreases with an increase in panel size. FST ranged from 0.182-0.649 with 10 SNVs to 0.393-0.468 with 500 SNVs (mean FST = 0.427) and the interquartile range (IQR) was reduced by 6.3-fold from 0.133 to 0.021 between these two datasets.

**Figure 4.**
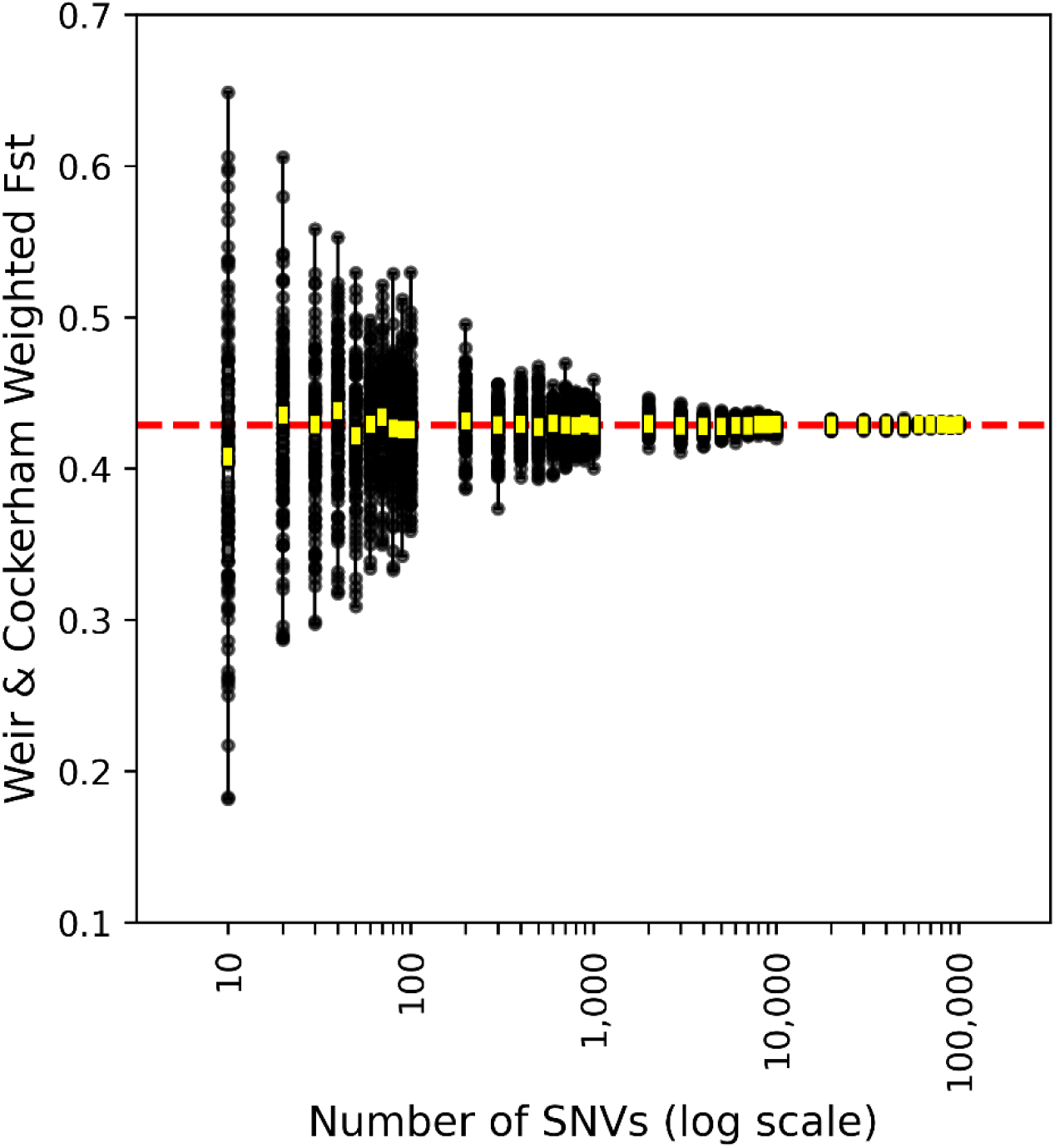
Fixation index (FST) versus panel size. The global, weighted Weir-Cockerham FST was measured between northern *S. haematobium*, southern *S. haematobium*, and *S. bovis* with panels ranging from 10-100,000 SNVs. The red line indicates FST measured with all autosomal, unlinked, common (minor allele frequency >0.05) SNVs in the whole genome dataset (n_SNVs_ = 381,501). Yellow boxes represent mean values for each panel size. Individual FST values are shown with black dots. Mean Fst is relatively consistent regardless of panel size, but the variation among replicates is inversely correlated the number of makers. The source data shown in this figure is available in the supplemental materials.

Supplemental Tables 1 and 2 provide estimated costs to genotype 1,000 samples with whole genome SNVs or a 500-locus amplicon panel. Sequencing an amplicon panel costs ∼$29.43 per sample compared $77.21 for whole genome sequencing. Of these costs, $9.34 goes towards isolating, purifying, and whole genome amplifying DNA from miracidia stored on FTA cards wich is used in both methods. Consumables like pipette tips, tubes, *etc.* count for an additional $2.16-$4.40 per sample. Library preparation for whole genome sequencing requires additional steps leading to higher consumable costs. Sequencing costs are higher for whole genome sequencing since each sample needs approximately three orders of magnitude more reads to achieve acceptable coverage for genotyping; 7.5Mb per amplicon sample vs 8.0 Gb per whole genome sample. Alternatively generating an initial pool of 100 uM primers for 500 loci represents a substantial upfront cost (∼$5,400) that offsets some of these savings when using amplicon panel. Pooled primers are sufficient for thousands of reactions so are ideal for large studies with multiple individuals.

## Discussion

Population genetics can be used to guide the management and elimination of schistosomes (41). Unfortunately, the most common tools used for studying schistosome population genetics and hybridization - *cox*1/ITS or whole genome sequencing - represent the extremes in costs, throughput, and precision. A two-marker system provides little information concerning ancestry in schistosomes. Further, *in-silico* work has shown that 7-293 species diagnostic markers are needed to differentiate between contemporary hybrids from parental species. However, differentiating advanced backcrosses can require thousands of markers (42, 43). In this study, we used *in-silico* tests to examine the performance of random SNV panels relative to whole genome data. The results show that panels comprising hundreds of SNVs contain comparable information to hundreds of thousands of markers (Figure 1, Table 1). This is similar to results from human studies that show 500 -1,000 random SNVs can effectively replicate whole genome results (44).

Generating sequencing libraries from hundreds of loci can be done in a multiplex PCR reaction. Target loci can be randomly selected from available variants, and primers pools that minimize mispriming and optimize PCR reaction conditions can be generated with tools like PrimerJinn (45), BatchPrimer3 (46), or SADDLE (47). After the target loci have been amplified a second PCR reaction is used to index the library and complete the Illumina sequencing adapters. This method has been used to genotype thousands of individuals at hundreds of loci in other taxa (26).

### Random SNVs vs Ancestry Informative Markers

Other studies have used tools like PCA loadings or FST to extract a minimal set of <u>a</u>ncestry informative <u>m</u>arkers (AIMs) to distinguish between species or populations (33, 48–50). These panels may contain as few as a dozen markers (51).

For studies where the central goal is to evaluate levels of hybridization between *S. haematobium* and *S. bovis*, such AIM panels may be useful. Modeling work has shown that identifying F1 hybrids requires at least 5 species diagnostic markers (52). If markers are not diagnostic, as many as 12 loci are needed to identify an F1 hybrid only if there are moderate allele frequency differences between the parental populations (53). That number climbs dramatically when trying to differentiate between advanced backcrossed from pure-parental classes. For example, after 6 generations of backcrossing, 191 diagnostic markers are required to achieve 95% confidence (α = .05) and 293 markers are needed for 99% (α = .01) confidence in the classification (42). There are 275,657 SNVs that differentiate *S. haematobium* and *S. bovis* (Supplemental Table 2 in 14): a subset of these could be used to generate an AIM panel. However, while AIM panels maximize the amount of information regarding ancestry or population assignment, they are biased when quantifying population genetic parameters. Hence, a small panel of AIMs would be useful to explore levels of introgression between *S. haematobium* and *S. bovis*, but the same panel would be extremely biased for studying population differentiation between locations, or determining relatedness of parasites, and may miss signals of hybridization or introgression from other species (20, 54–56). Further, our results show that small panels of SNVs may lack precision in estimating parameters like FST (Figure 4). Both random and AIM panels may be useful for different research questions, and optimal panels for both can be designed based on genome-wide marker data from Platt *et al*. (14). However, randomly selected marker panels are more generally useful.

### Cost-efficiency vs Diagnostic Power

The accuracy of hybrid identification or population parameter estimation is directly linked to the number of markers used, but additional markers increase costs. We estimated the costs to examine 1,000 parasitesusing a 500-locus amplicon panel and whole genome sequencing. The results show that amplicon panels are two to three times more affordable than whole genome sequencing while providing comparable resolution. Further technical modification can decrease the prices even further. For example, cost-saving techniques like library miniaturization (57) and automation can substantially reduce the price of reaction sizes and the necessary reagents. Other reduced representation methods likely provide similar benefits. For example, Landeryou *et al*. identified *S. haematobium* and *S. guineensis* hybrids by genotyping RADseq loci (20). The costs of RADseq are comparable to amplicon panels given the reduced number of reads needed to achieve reasonable coverage. However, amplicon panels require less technical expertise than RADseq, so they can be widely adopted. Laboratories that routinely conduct *cox*1/ITS genotyping can transition to using amplicon panels. In addition to being cost-effective, amplicon panels require fewer computational resources for data analyses. For example, whole genome SNV datasets with millions of markers require access to a computing cluster. Our lab uses a computing cluster with 12 nodes; each with 192 processors and 1-1.5 Tb of memory. In that case, genotyping 218 samples still took several weeks. By comparison, smaller amplicon datasets could be analyzed on a reasonably powered personal computer in a matter of hours.

### Limitations and Future Perspectives

While reduced representation sequencing seems to be a viable option for schistosomes it is worth noting a few limitations. First, while the median difference between *S. bovis* ancestry estimates from genome sequence data and an amplicon panel of 500 markers is less than 1%, individual estimates may differ by -10.3% to 5%. This variance could affect hybrid classification with software that uses allele frequencies (43). Future work could incorporate lab generated hybrids as a positive control to precisely quantify this uncertainty. Second, the SNVs in this study are circulating in the *S. haematobium* and *S. bovis* populations at moderate frequencies. Introgression from an unsampled species, like *S. guineensis* (20), may not be readily apparent in this case. They could even be missed if the introgressed variants are at low frequencies (<5%) and were filtered out from the Platt *et al*. (14) SNV set. This issue will be resolved as the population genomic data for *S. curassoni, intercalatum*, *guineensis*, *mattheei* and *margrebowiei* become available and panels can be re- designed and updated to have the broadest applicability within the *S. haematobium* group.

## Conclusion

Large scale, genome studies are able to resolve population genetic structure with high precision but are difficult to scale adequately to detect rare hybridization events. By contrast, genotyping parasites with only two markers, as is the standard in many schistosome studies, can be applied to thousands of individuals, but can produce ambiguous results. Here we show that sequencing a moderate number of random SNVs can efficiently capture the majority of the population genetic data contained within a parasite genome. Further, the costs of sequencing such panels are considerably less than whole genome sequencing allowing examination of hundreds or thousands of individuals. We conclude that amplicon panels provide an effective compromise strategy, allowing cost-effective, robust characterization of large numbers of samples, and providing accurate measures of both population structure and hybrid ancestry.

## Acknowledgements

We thank Sandy Smith and John Heaner for providing computational support at Texas Biomedical Research Institute’s High-Performance Computing Cluster. Jenna M. Hulke provided valuable insights regarding data analysis and experimental design. This research was funded by the National Institute of Allergy and Infectious Diseases (NIAD R01 AI166049-01).

